# SARS-CoV-2 Main Protease: a Kinetic Approach

**DOI:** 10.1101/2022.05.01.490203

**Authors:** Thierry Rebetez

## Abstract

In this article, I present a new model of the interaction of the main protease (M^pro^) from SARS-CoV-2 virus with its substrate. The reaction scheme used to describe this mechanism is an extension of the well-known Michaelis-Menten model proposed in 1913 by Leonor Michaelis and Maud Menten [1]. The model I present here takes into account that one M^pro^ enzyme monomer interacts with another M^pro^ monomer in the presence of the substrate, leading to the formation of an enzyme dimer bound to one substrate molecule. Indeed, this dimer is formed by the sequentially binding of one M^pro^ enzyme monomer to one molecule of substrate, followed by another M^pro^ enzyme monomer binding to this M^pro^-substrate complex. This reaction mechanism is also known in the literature as substrate-induced dimerization [3]. Starting from this new reaction scheme established for this catalytic mechanism, I derived a mathematical expression describing the catalytic rate of the active M^pro^ enzyme dimer as a function of the substrate concentration [*S*]. The plot corresponding to this substrate-induced dimerization reaction shows a function *f* ([*S*]) that is not monotonic, *i*.*e*. not strictly increasing or decreasing, but with a second derivative initially negative and then becoming positive after having passed the *V*_*max*_ point. This is typically a type of curve showing a phenomenon like the one of substrate inhibition (for instance, inhibition by excess-substrate [7]). The graphical representation of this process shows an interesting behaviour: from zero *μ*M/s, the reaction rate increases progressively, similar to the kind of curve described by the Michaelis-Menten model. However, after having reached its maximum catalytic rate, *V*_*max*_, the reaction rate decreases progressively as we continue to increase the substrate concentration. I propose an explanation to this interesting behavior. At the moment where *V*_*cat*_ is maximum, we can assume that, in theory, every single substrate molecule in solution is bound to two enzyme monomers (*i*.*e*. to one active dimer). The catalytic rate is thus theoretically maximized. At the time where the reaction rate begins to decrease, we observe a new phenomenon that appears: the enzyme monomers begin to be “diluted” in the solution containing the excess substrate. The dimers begin to dissociate and to bind increasingly to the substrate as inactive monomers instead of active dimers. Hence, it is more and more unlikely for the enzyme monomers to sequentially bind twice to the same substrate molecule (here, [*E*] *≪* [*S*]). Thus, at this stage, the substrate-induced dimerization occurs less often. At the limit, when the substrate is in high excess, there is virtually no more dimerization which occurs. This is one example of excess-substrate inhibition. Furthermore, after having established this fact, I wanted to see if this catalytic behavior was also observed *in vitro*. Therefore, I conducted an experiment where I measured the catalytic rate of the M^pro^ dimer for different substrate concentrations. The properties of my substrate construct were such, that I could determine the catalytic rate of the enzyme dimer by directly measuring the spectrophotometric absorbance of the cleaved substrate at *λ* = 405 nm. The results show explicitly — within a margin of error — that the overall shape of the experimental curve looks like the one of the theoretical curve. I thus conclude that the biochemical behavior of the M^pro^ *in vitro* follows a new path when it is in contact with its substrate: an excess substrate concentration decreases the activity of the enzyme by the phenomenon of a type of excess-substrate inhibition. This finding could open a new door in the discovery of drugs directed against the M^pro^ enzyme of the SARS-CoV-2 virus, acting on the inhibition by excess-substrate of the M^pro^ enzyme, this protein being a key component in the metabolism of the virus. Furthermore, I have established that the maximum of the fitted curve, *V*_*max*_, depends only on [*E*]_*T*_ and not on [*S*]. 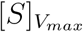 exhibits the same dependence pattern. Therefore, if I keep [*E*]_*T*_ close to zero, the catalytic rate of the enzyme will also be greatly reduced, which can be understood intuitively. Finally, if we dilute the enzyme sufficiently in the host cell by injecting a suitably high concentration of the octapeptide substrate AVLQSGFR (an inhibitor of the original substrate), this artificial substrate will bind to the “intermediate” dimer from the polypeptide and prevent the precursor M^pro^ from auto-cleaving and dimerizing due to the “distorted key” effect of the octapeptide on the “intermediate” dimer. The precursor peptide M^pro^ will auto-cleave to a lesser extent than in the absence of the artificial octapeptide and thus the concentration of the total enzyme [*E*]_*T*_ will be lowered in the cell. It would therefore be possible to control the virulence of the virus by adjusting the concentration of the artificial inhibitory octapeptide. However, this is only speculation and has yet to be verified in practice.

## 1 Introduction

The coronavirus disease 2019, or Covid-19, is an infectious disease caused by the virus SARS-CoV-2 (Severe Acute Respiratory Syndrome CoronaVirus 2). Its most serious symptoms are an acute respiratory distress syndrome (ARDS) [8] or a cytokine release syndrome (CRS) [9]. Human-to-human transmission is mainly by respiratory droplets and aerosolization [10]. In the early 2020s, the coronavirus 2019 is a pandemic, which disrupted human activity across the entire planet, through general confinements, strict sanitary measures, border closures, and the slowdown or cancellation in many economic sectors or events [11]. Several types of anti-covid vaccines are being manufactured, including messenger RNA vaccines [12][13] which, starting in 2021, are administered to a large part of the population, more widely in rich countries, in Europe, North and South America, Asia and Oceania [14], in an attempt to curb the pandemic. In September 2021, the World Health Organization (WHO) offered mRNA technology to Bangladesh, Indonesia, Pakistan, Serbia and Vietnam in order to increase manufacturing of Covid-19 vaccines for poorer countries [15]. The struggle is going on…

One key enzyme in the metabolism of the SARS-CoV-2 virus is M^pro^ (also named 3CL^pro^, or 3C-like protease for 3-chymotrypsin-like cysteine protease). M^pro^ is an ideal target for antiviral drug design due to its high conservation between different coronavirus strains and absence of functional analogs in the human proteome [23]. It is noteworthy that M^pro^ from SARS-CoV-1 and SARS-CoV-2 are structurally and functionally similar (their RNA genomes are about 82% identical) and also similar to M^pro^ from MERS (Middle East Respiratory Syndrome) [24][25].

M^pro^ consists of a polypeptide chain of 306 amino acids comprising 3 domains per monomer. Structurally, two monomers orient perpendicular to one another to form a dimer. The substrate-binding pocket is located in a cleft between domains I and II and the active site consists of a Cys145–His41 catalytic dyad [26]. It is a cysteine protease that plays a crucial role in the virus’ life cycle. Hence, after infection, the virus releases two overlapping replicases, polyproteins pp1a and pp1ab, that are functional polypeptides and essential for the replication and transcription of the virus [27]. Two viral proteases, M^pro^ (main protease, Nsp5) and PL^pro^ (papain-like protease, Nsp3), cleave polyproteins 1a and 1ab. M^pro^ is first autocleaved from polyproteins to yield the mature enzyme. Further, the proteolytic cleavage by M^pro^ corresponds to 13 non-structural protein (Nsp4 to Nsp16), while PL^pro^ cleaves three peptide bonds, liberating three proteins (Nsp1 to Nsp3) [28]. Each Nsp has a specific role in the life cycle and pathogenicity of the virus [29]. We see that M^pro^ has a pivotal role in the metabolism of the SARS-CoV-2, and the cleavage of these 16 Nsp by M^pro^ and PL^pro^ is crucial to the virulence of the virus. M^pro^ is thus a potential target to antiviral agents acting on its central role in the metabolism of the SARS-CoV-2.

The enzyme exists as a mixture of dimers and monomers, and only the dimers are considered to be active. Both the N-and C-terminus of M^pro^ have been shown to be critical for dimer formation and for enzyme function [16]. Each subunit has its own substrate binding site located at the interface between domains I and II. Furthermore, domain III is involved in the dimerization of M^pro^ [17]. Nevertheless, only one active protomer in the dimer of M^pro^ is enough for catalysis [3]. [18] showed that the monomers are always inactive, that the two protomers in the dimer are asymmetric, and that only one protomer is active at a time. [2] showed that residue Glu-166 of M^pro^ plays a pivotal role in connecting the substrate binding site with the dimer interface. Substrate binding to a monomer induces a conformational change of the enzyme that shifts the equilibrium from monomer to the catalytic-competent dimer. This mechanism is called substrate-induced dimerization. Then, catalysis and release of product, followed by dissociation and regeneration of the inactive protomers, complete the catalytic cycle. This mechanism is now well accepted in the literature and is compatible with other studies ([3], [4], [5]). For instance, Ho et al. ([21]) have shown that the dual role of the conserved Glu residue (Glu-166 in SARS-CoV-2 M^pro^ and Glu-169 in MERS-CoV M^pro^) in catalysis and dimerization is consistent for both M^pro^s. Lai et al. [20] reported that SARS-CoV-2 M^pro^ exists as a monomer/dimer mixture at a relatively high protein concentration in solution (4 mg/mL) and is exclusively monomeric at a lower protein concentration (0.2 mg/mL) in solution, and only the dimer was the active form of the proteinase. In my study, I have thus chosen an initial guessed enzyme concentration of 0.5 *μ*M (33.8 *μ*g/mL) for my catalytic rate determination experiment. This guess turned out to be valid for the purpose of the assay: indeed, one of the criteria that must be fulfilled in order to have a valid model is following: the majority of the dimers in solution must come from substrate-induced dimerization and not from spontaneous dimerization in solution. This assumption is synonym to say that the enzyme must dimerize only in the presence of the substrate.

## 2 Materials and Methods

### 2.1 Catalytic assay

The colorimetry-based peptide substrate, TSAVLQ-para-nitroanilide (TQ6-pNA) (Sigma-Aldrich Chemie GmbH, Industriestrasse 25, 9471 Buchs, Switzerland), was used to measure the proteolytic activity of SARS-CoV-2 M^pro^ throughout the course of the study as described previously [2], [19]. This substrate is cleaved at the Gln-pNA bond to release free pNA, resulting in an increase in absorbance at 405 nm. This absorbance was monitored using an Amersham Pharmacia Ultrospec 3100 Pro UV/VIS spectrophotometer. The protease activity assay was performed in 25 mM HEPES buffer, 0.2% Tween-20, pH 7.0 at 25 °C. The substrate stock solution was 2650 *μ*M and the working concentrations were from 0 to 1200 *μ*M. In the substrate catalytic assay, the concentration of SARS-CoV-2 M^pro^ was of 0.5 *μ*M (33.8 *μ*g/mL). The derivation of the catalytic rate of M^pro^ was done using the Beer-Lambert law, with a molar extinction coefficient of 9.96E-3 *μ*M^*−*1^cm^*−*1^ at 405 nm, a reaction time of 240 s per cuvette and a cuvette depth of 1 cm.

### 2.2 Determination of the rate constants by the nonlinear least squares method

*A basic problem in science is to fit a model to observations subject to errors. It is clear that the more observations that are available, the more accurately will it be possible to calculate the parameters in the model. This gives rise to the problem of “solving” an overdetermined linear or nonlinear system of equations. It can be shown that the solution which minimizes a weighted sum of the squares of the residual is optimal in a certain sense*.

Åke Björck, Numerical Methods for Least Squares Problems, SIAM, 1996.

Quasi-steady state enzyme kinetic parameters (*k*_1_ to *k*_5_ in our case) were obtained by fitting the velocity data (different *V*_*cat*_ and [*S*] measured during the assay) to the model equations (eqs. (19) to (24)) in MATLAB^®^. I used the function

`lsqcurvefit`

which enabled me to fit a parameterized nonlinear function to data. I used the following initial guess for the parameters to fit the model to the data:

`x0 = [10, 15, 15, 15, 15]`

## 3 Results

Let us define the following concentrations:

[*S*] := Concentration of the substrate molecules unbound.

[*E*] := Concentration of the enzyme monomers unbound.

[*ES*] := Concentration of the substrate molecules bound to one enzyme monomer.

[*E*_2_*S*] := Concentration of the substrate molecules bound to two enzyme monomers (= to one enzyme homodimer).

[*P*] := Concentration of the product.

Reactional scheme:

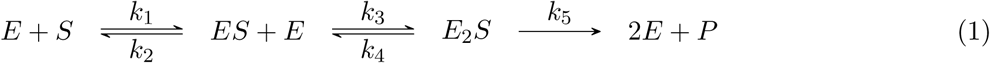

Evolution equations:

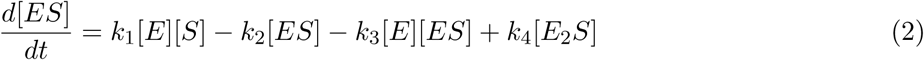

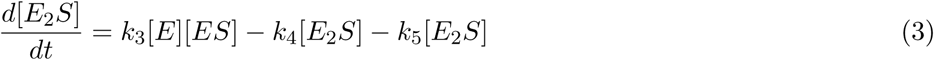

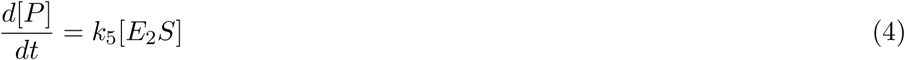

Quasi-steady state assumption (QSSA):

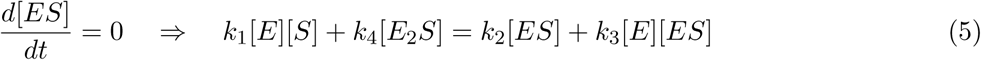

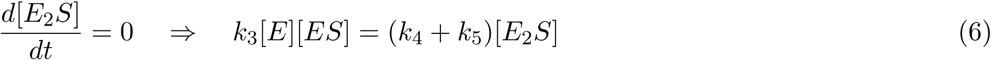

From these 2 equations, we find:

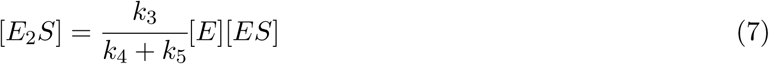

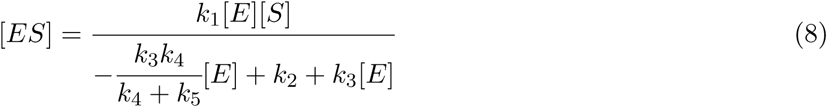

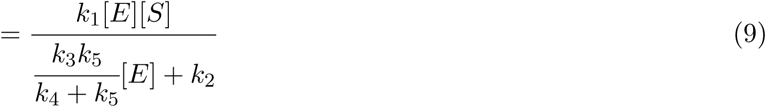

Let:

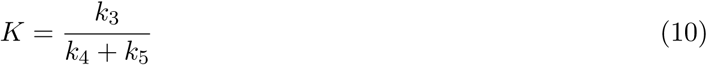

Then:

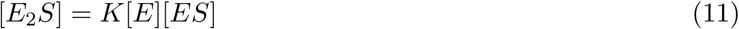

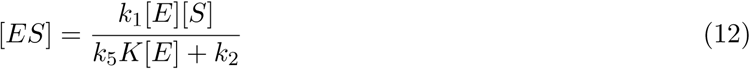

The rate of production of P is:

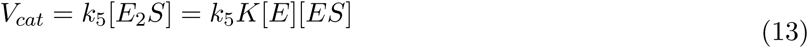

We want to express [*E*_2_*S*] as a function of [*E*]_*T*_ and [*S*].

Conservation of the enzyme:

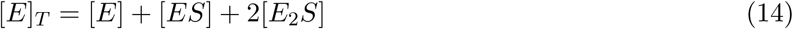

We can combine eqs. (11), (12) and (14):

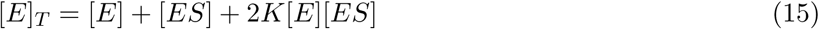

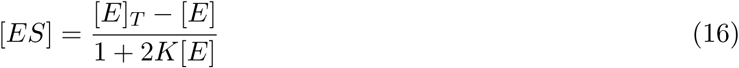

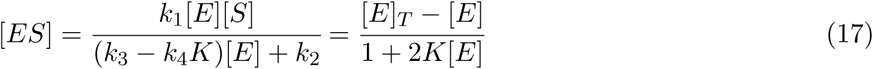

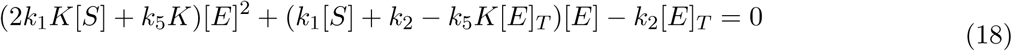

The solutions of this quadratic equation are:

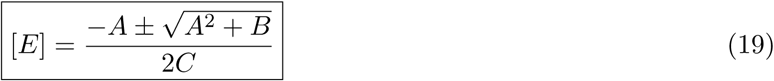

With

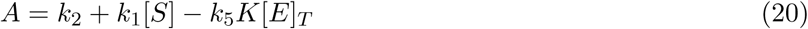

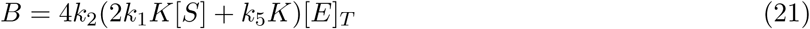

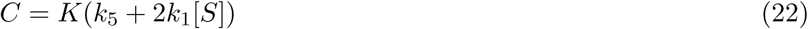

N. B.: To get [*E*] *>* 0, we need to take the solution with “+”.

The rate of production of the product P, 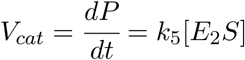, can thus be expressed as:

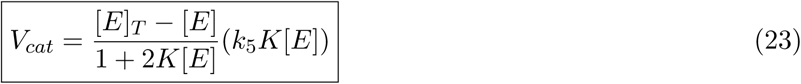

Or, expressed slightly differently:

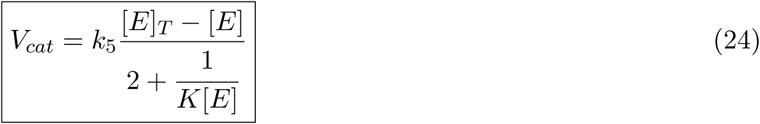

with [E] given by eq. (19), and

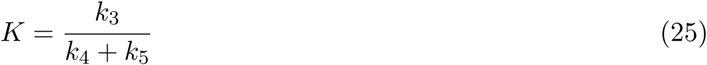

## 4 Discussion

### 4.1 Comparison of theory with the *in vitro* experiment

The graph of Fig. 1 shows the theoretical behaviour of *V*_*cat*_ according to the substrate concentration. We can see that for every set of rate parameters (red, black and green curves, see also Table 1), the curves show a similar non monotonic shape, with an increase until a maximum, and after having reached this peak, the catalytic rate decreases progressively towards the asymptote *y* = 0 at different pace according to the rate parameters. I propose the following explanation to this behaviour: the first part of the curve — *i*.*e*. when the enzyme catalytic rate follows a quasi Michaelis-Menten typical curve scheme until *V*_*max*_ is reached — corresponds to the state where the substrate concentration in solution is relatively low: the concentration of the enzyme compared to [*S*] is sufficiently large to allow a dimerization of the enzyme monomers induced by the substrate (see also Fig. 2). We have here the case where two enzyme monomers are bound to one substrate and form a substrate-bound dimer (*E*_2_*S*). Then, after having reached 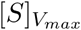 where the maximum velocity *V*_*max*_ is reached, the concentration of the substrate is increased more and more, leading to a concentration range where the enzyme monomers are so “diluted” in the solution of excess substrate that it is almost unlikely (from a statistical point of view) for the enzyme monomers to get close enough to each other to enter into chemical interaction via a substrate molecule: they are not able to bind sequentially to a single substrate molecule anymore as it is the case in the dimerization event. In this scenario, we have [*E*] *≪* [*S*].

**Table 1:**
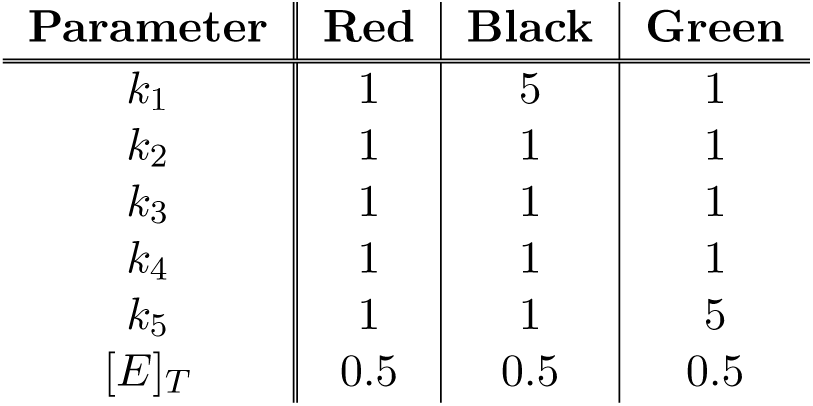
Parameter values and total enzyme concentration in the substrate-induced dimerization model. This choice of distinct parameter values and of total enzyme concentration results in different theoretical curves for *V*_*cat*_. These curves have nevertheless a very similar aspect each other. **W**e observe that all three red, black and green curves in Fig. 1 tend to zero when [*S*] tends towards infinity, but more or less quickly according to the curve and to its different rate constants parameters. For the different units of the rate constants, see section 4.2 Dimensional Analysis below.

| Parameter | Red | Black | Green |
| --- | --- | --- | --- |
| $k_1$ | 1 | 5 | 1 |
| $k_2$ | 1 | 1 | 1 |
| $k_3$ | 1 | 1 | 1 |
| $k_4$ | 1 | 1 | 1 |
| $k_5$ | 1 | 1 | 5 |
| $[E]_T$ | 0.5 | 0.5 | 0.5 |

**Figure 1:**
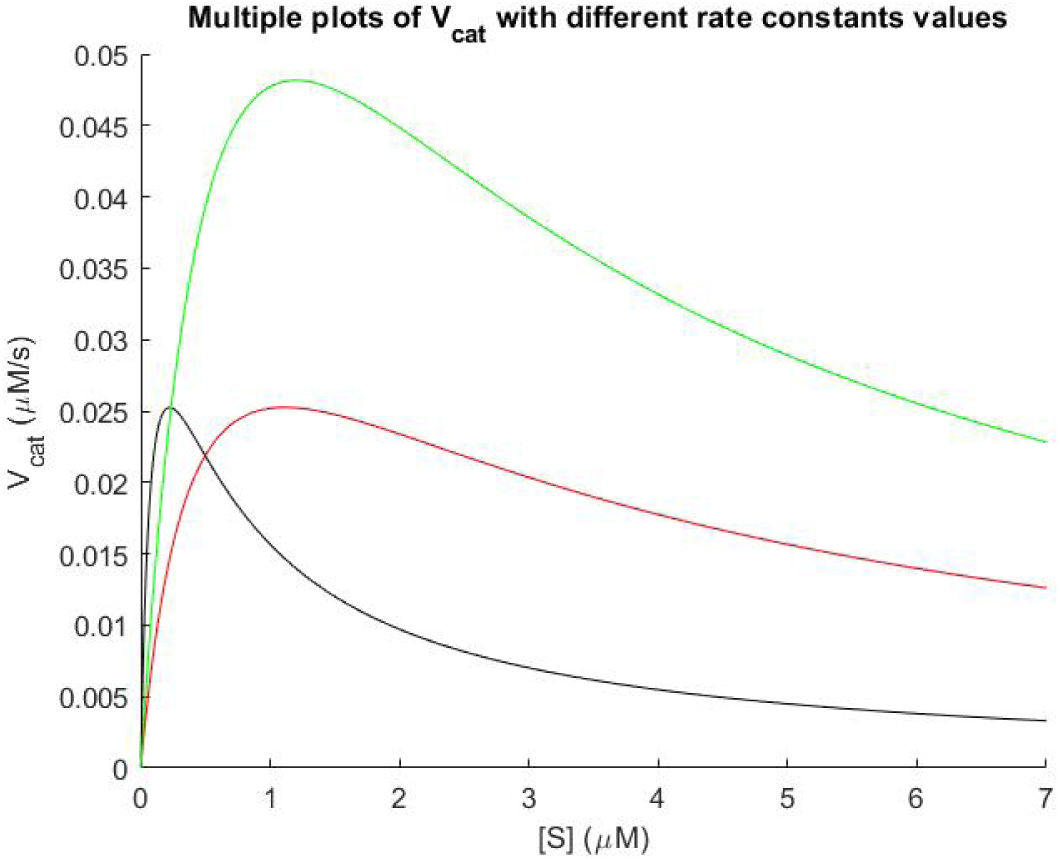
Rate of production of P (*V*_*cat*_) in function of the substrate concentration [*S*] for different parameter values (see Table 1). Overview of the theoretical principle.

**Figure 2:**
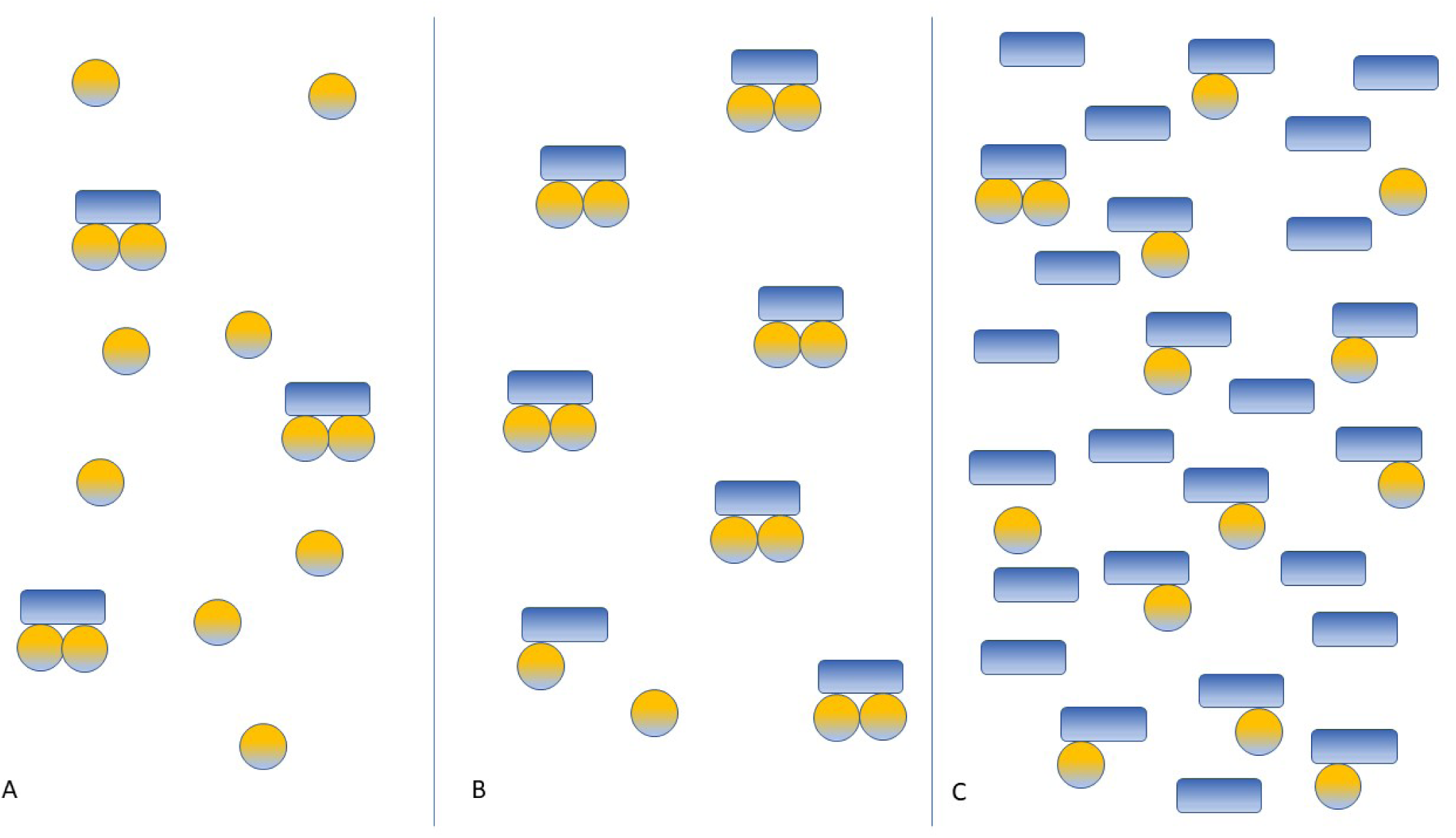
Schematic illustration of the behavior of interactions between the enzyme M^pro^ (yellow spheres) and the substrate S (blue rectangles).

At part A of Fig. 2, we see that there is an excess of enzymes (monomers, represented as yellow spheres) and that the substrate is underrepresented. This state thus corresponds to the part of Fig. 1 where the graph has a hyperbolic appearance (part of the graph to the left of 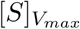). In this case, almost each substrate molecule is bound to two monomers of the enzyme: the substrate is saturated with enzyme dimers.

At part B of the illustration, we have the case where the concentrations of the substrate and the enzyme are optimally weighted with respect to each other: the monomers of the enzyme saturate the substrate. We have reached *V*_*max*_ and almost every substrate has caused dimerization of the enzyme through the substrate-induced dimerization mechanism. At this point, the catalytic rate has reached a maximum.

On the right side of the diagram (part C), we see what happens when we add even more substrate molecules to the solution. Having passed beyond the maximum catalytic rate (*V*_*max*_), the monomers dissociate gradually from the substrate (according to their dynamic equilibrium) and return to solution. At this point of the system, the monomers are statistically and progressively diluted in the substrate solution when more and more substrate is added to the solution. Therefore, the following case is gradually becoming preponderant: an enzyme monomer binds to a substrate molecule but the substrate-induced dimerization process does not occur any more. On the contrary, the enzyme monomers are “distributed” and “diluted” throughout the almost entire solution where there is an excess of substrate. Statistically, we observe almost exclusively one substrate molecule bound to one enzyme monomer, if any. Thus, the catalytic rate decreases progressively when we add more and more substrate to the solution. In this scenario, *V*_*cat*_ progressively and asymptotically approaches the x-axis, namely *y* = 0, when we are dealing with huge amounts of *S*. Eventually, with [*S*] sufficiently high, *i*.*e*. when [S] approaches infinity, *V*_*cat*_ tends towards zero and vanishes completely.

### 4.2 Dimensional analysis

In order to know and to verify the dimensions of the units of the different rate constants, we performed a dimensional analysis. We get the following results:

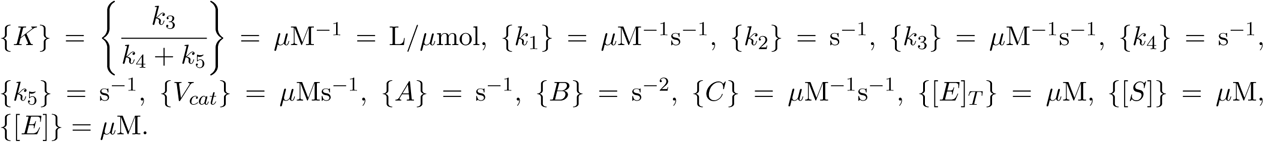

### 4.3 Determination of the rate constants by the nonlinear least squares method

After computation in MATLAB^®^, I get the nonlinear least squares solution shown in Fig. 3.

**Figure 3:**
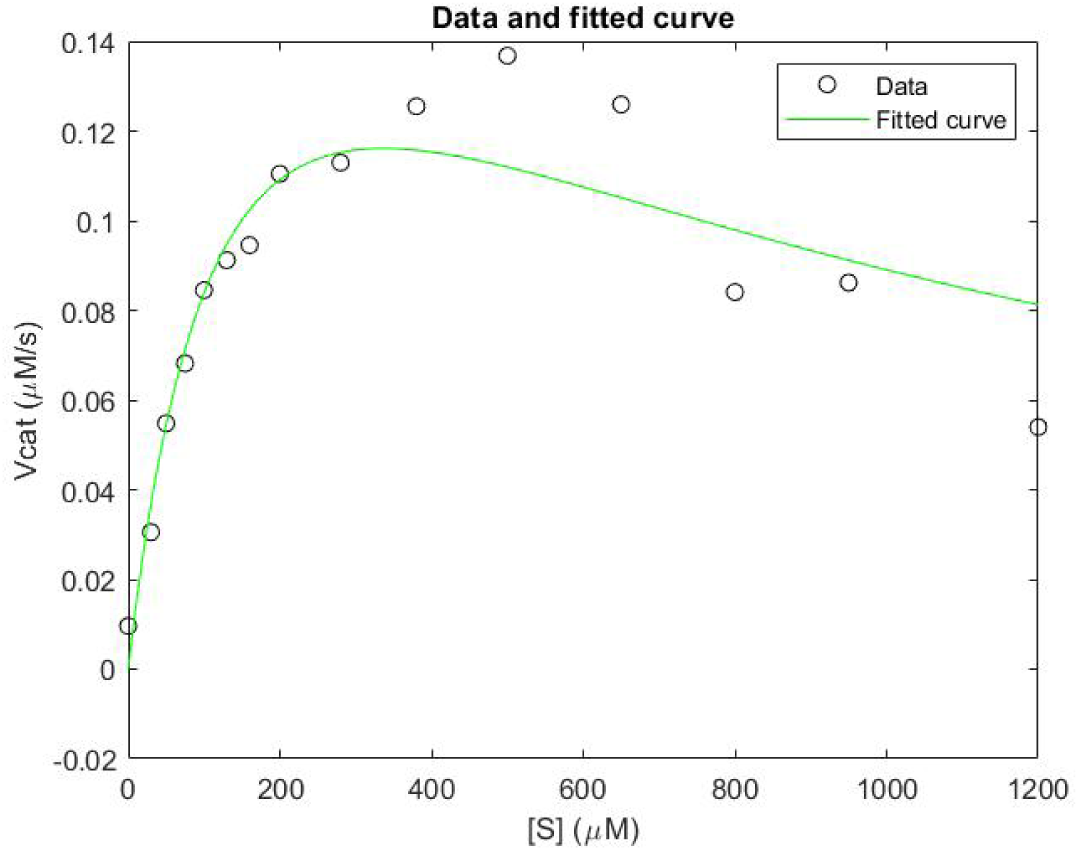
Fitting of the non-linear parameterized function to experimentally obtained data.

On this graph, MATLAB^®^ has chosen the parameters such that the distance between the data and the curve is minimal, *i*.*e*. the curve that fits best. In this case, the fit is good, but not perfect. It will therefore be necessary to repeat the experiment several times to verify the reproducibility of the system. Nevertheless, we can use this data later in this demonstration. This fit gives the following parameter values:

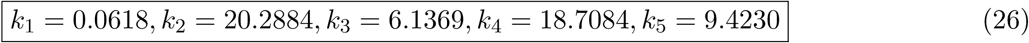

The units of the different k’s of box (26) are given in section 4.2 Dimensional analysis.

### 4.4 Determination of *V*_*max*_ and of 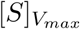

We have:

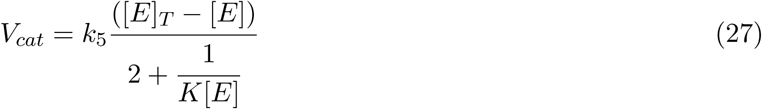

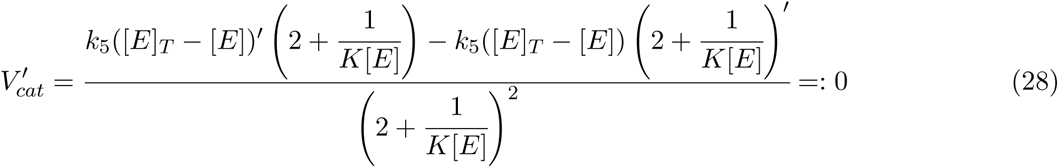

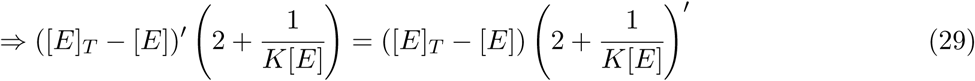

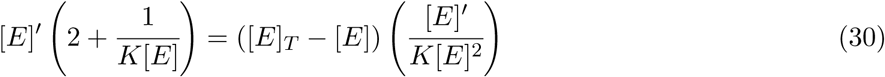

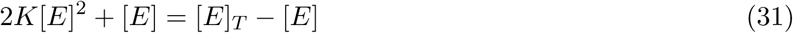

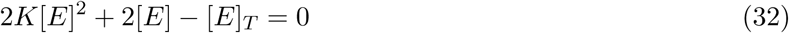

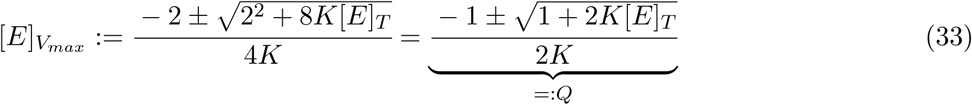

at the point 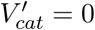. To get [*E*] *>* 0, we need to take the solution with “ + “.

If we put this last result (eq. (33)) in eq. (27) and do some algebra, we get an expression for *V*_*max*_:

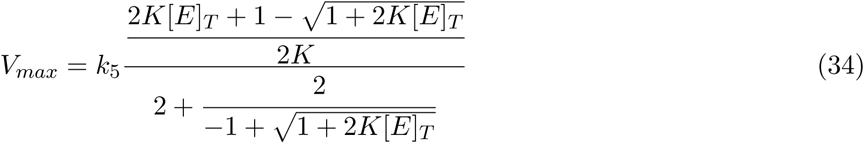

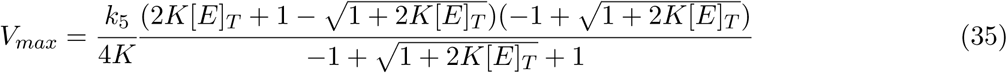

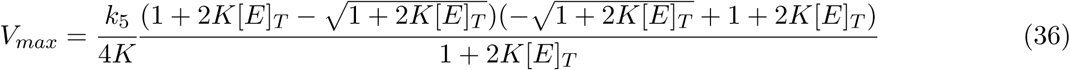

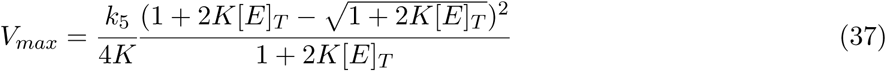

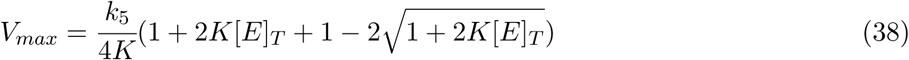

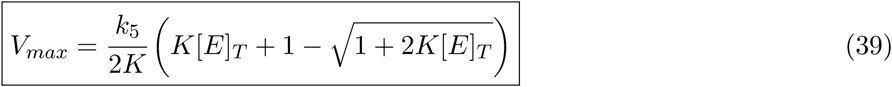

We can see that this theoretical calculation of *V*_*max*_ (eq. (39)) agrees numerically with the fitted curve (see 2^*nd*^ row of Table 2 below corresponding to [*E*]_*T*_ = 0.5 *μ*M).

Moreover, if we take the eq. (19) and make it equal to *Q* (eq. (33)), we get the following equality:

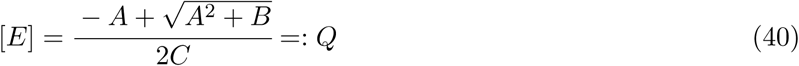

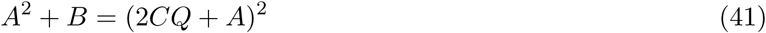

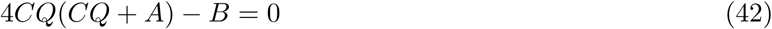

If we replace the values of A, B and C (eqs. (20), (21) and (22)), and the value of Q (eq. (33) in this last equation, we get the following expression:

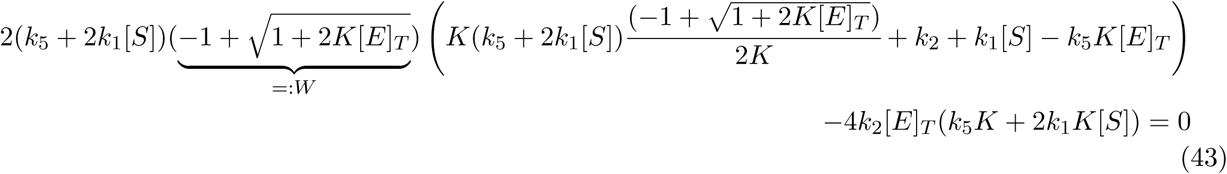

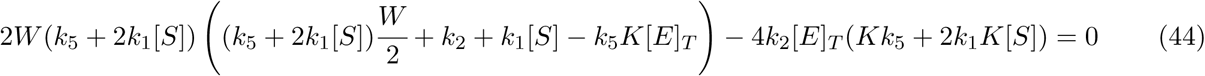

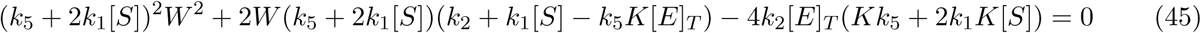

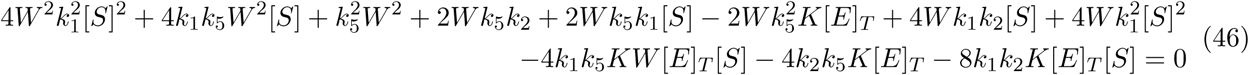

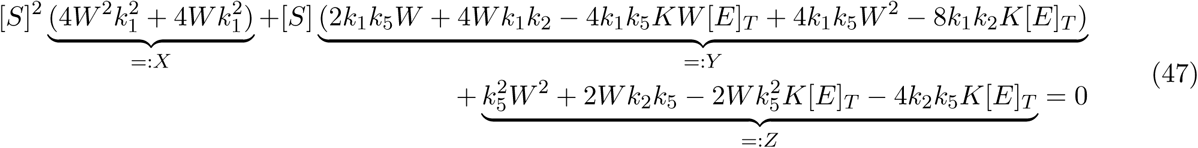

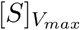 is thus equal to:

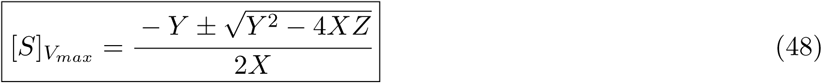

with X, Y and Z given above in eq. (47) and W in eq. (43).

Here, *W >* 0, *X >* 0. We have found here the numerical value of 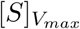 only by calculation. We can easily verify that, if we take the values mentioned above for the rate constants (chap. 4.3), inject them into (48), choose the arbitrary value of 0.5 *μ*M for [*E*]_*T*_ (like the one for the experimental data, see chap. 2 Material and Methods), we indeed find that the numerical value of 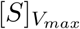 (in this case 336.1844 *μ*M) and *V*_*max*_ (0.1161 *μ*M/s) are in very good agreement with the value of 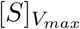 and *V*_*max*_ sampled on the fitted curve. We also see that *V*_*max*_ and 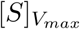 depend only on the value of [*E*]_*T*_ and the five rate constants. So I wanted to check the behavior of the point 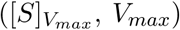 for different values of [*E*]_*T*_. The result is presented in Table 2 and in Fig. 4 and 5.

**Figure 4:**
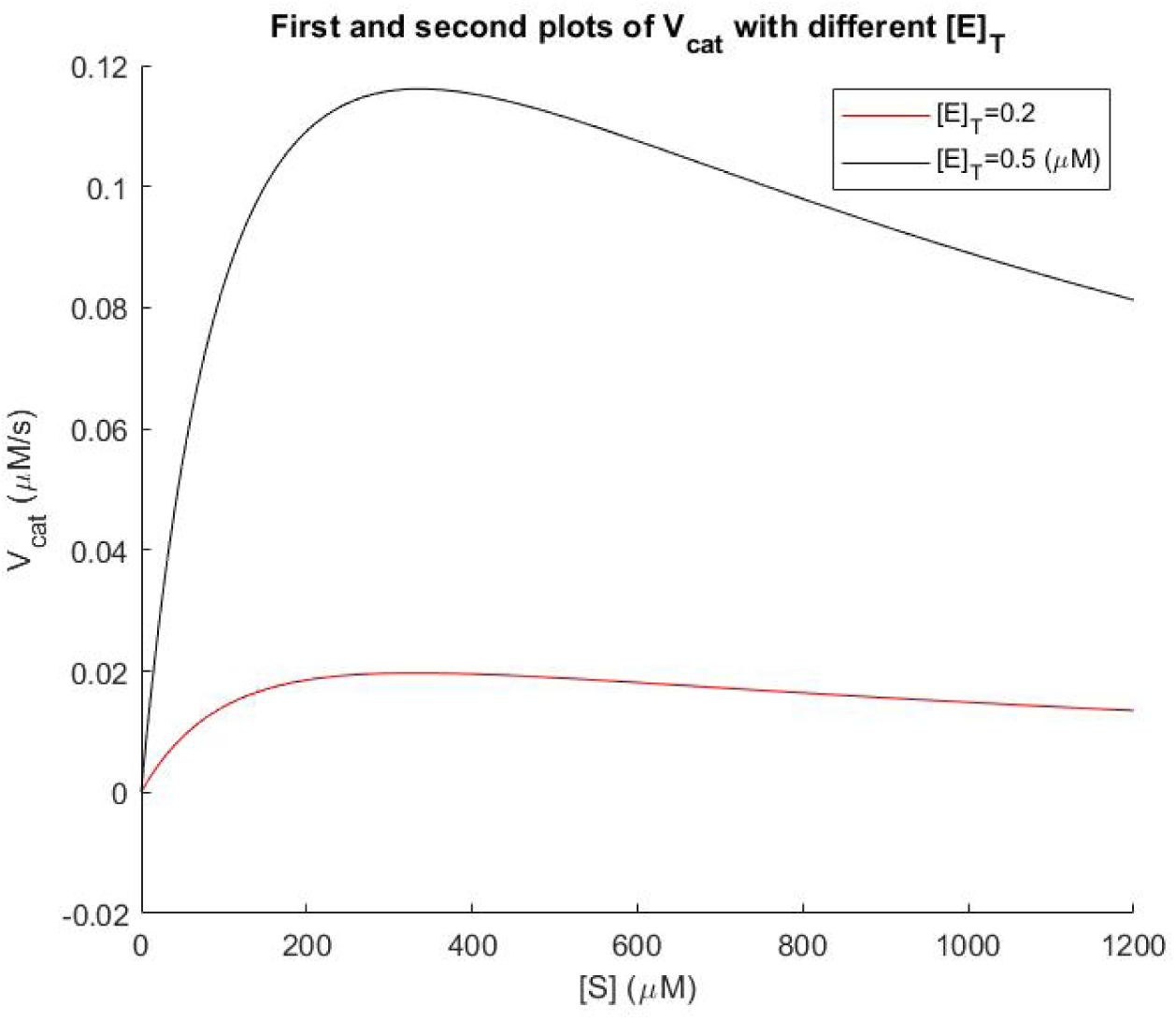
The maximum of the curve (point where the monotony of the curve is broken) is different for different values of [*E*]_*T*_.

**Figure 5:**
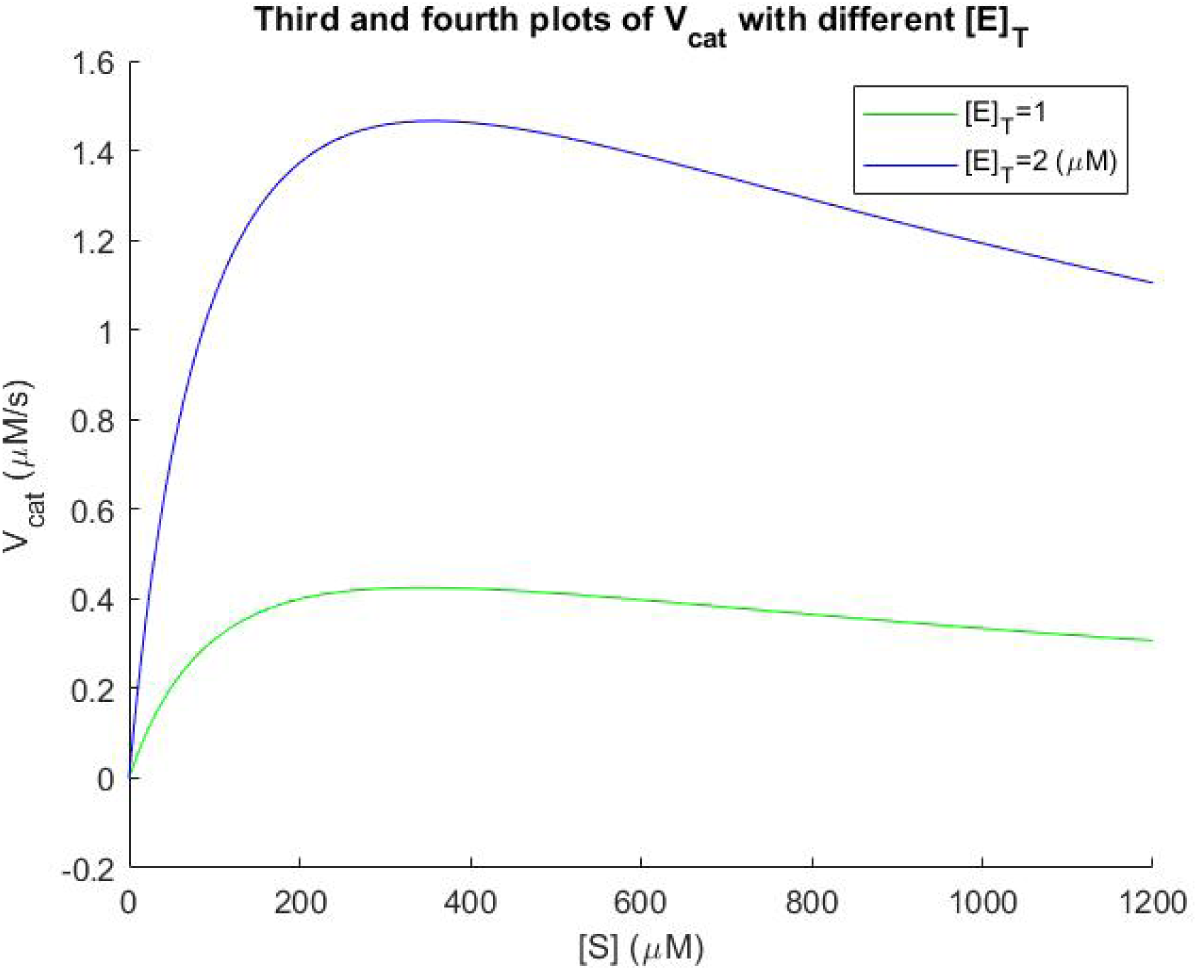
The maximum of the curve is different for different values of [*E*]_*T*_.

**Table 2:** For each of the four values of [*E*]_*T*_, I have computed the coordinates of the point 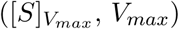 (see bold values in Table 2). These numbers are purely numerical and theoretical values derived from eqs. (39) and (48). If we compare them to the actual coordinates of the peaks in the four different graphs in Figs. 4 and 5, which are practical data (derived from the five fitted rate constants), we see that theory (Table 2) and practice (Figs. 4 and 5) match almost perfectly! Our model perfectly describes the *in vitro* substrate-induced dimerization process. We can thus predict the value of the point coordinates 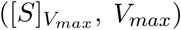 knowing *a priori* only the 5 rate constants (which are inherent to the enzymatic system and are themselves dependent only on temperature) and the total enzymatic concentration of M^pro^ ([*E*]_*T*_, see eq. (14)). It is enough to put these values into eqs. (39) and (48) and we then know the value of [*S*] from which *V*_*cat*_ starts to decrease, *i*.*e*. from which the virulence of the virus starts to reduce.

| $[E]_T$ | W | X | Y | Z | $[S]_{V_{max}} (\mu\text{M})$ | $V_{max} (\mu\text{M/s})$ |
| --- | --- | --- | --- | --- | --- | --- |
| 0.2 | 0.0427 | 6.8020E-4 | -0.1738 | -17.2067 | **331.7621** | **0.0197** |
| 0.5 | 0.1037 | 1.7485E-3 | -0.4545 | -44.8151 | **336.1844** | **0.1161** |
| 1 | 0.1984 | 3.6323E-3 | -0.9712 | -95.1532 | **343.6162** | **0.4253** |
| 2 | 0.3684 | 7.7014E-3 | -2.1579 | -209.2765 | **356.4340** | **1.4658** |

We can see that with increasing values of [*E*]_*T*_, the peak of the curve moves upwards and slightly to the right. These two last graphs show us that the value of the only parameter we can vary (*i*.*e*. [*E*]_*T*_) has a considerable impact on *V*_*max*_ of M^pro^, and consequently on the virulence of the virus. If we manage to keep [*E*]_*T*_ low, the top of the curve (*i*.*e. V*_*max*_) will shift downward and thus only a smaller percentage of M^pro^ will be active in this case. Thus, the final question to be addressed in this study is how to maintain a low concentration of total M^pro^ enzyme *in vivo*?

I will begin to answer this question by considering the octapeptide AVLQSGFR. In this paragraph we assume that SARS-CoV M^pro^ and SARS-CoV-2 M^pro^ share a quasi identical three-dimensional structure [30]. The prerequisite condition for a peptide to be cleaved by M^pro^ is a good fit and binding between the substrate and the enzyme’s active site. The octapeptide is a synthetic substrate for SARS-CoV M^pro^. Its sequence is derived from residues P4–P5’ of the N-terminal autoprocessing site of SARS-CoV M^pro^ and was identified by a docking study [31].

According to the “lock-and-key” mechanism in enzymology, this substrate binds to the active site of SARS-CoV M^pro^ and acts as a competitive inhibitor because of its “hybrid peptide bond” located between the subsites P1 and P1’. After chemical modification, this strong peptide bond is difficult to cleave. This mechanism illustrate the “distorted key” theory, where the “distorted key” can be inserted into a lock but can neither open the lock nor automatically get out from it [32]. That is why a molecule modified from a cleavable peptide can spontaneously become a competitive inhibitor against the enzyme [33].

Experiments show that the octapeptide AVLQSGFR is bound to SARS-CoV M^pro^ by six hydrogen bonds [31]. It is an effective inhibitor of SARS-CoV with an *EC*_50_ (half maximum effective concentration) of 2.7*·*10^*−*2^ mg/L and is capable of blocking virus replication. Furthermore, this octapeptide has no detectable toxicity in host cells.

Now let’s perform another thought experiment. Suppose we could inject a very large amount of this octapeptide substrate AVLQSGFR in the host’s cell. As it is very similar chemically and structurally to the original substrates of M^pro^ (*i*.*e*. the different subsites of the polyprotein to cleave in *trans*), it will shift, in our thought experiment, the equilibrium of the bounded M^pro^ towards this synthetic substrate, and the original substrates will have less M^pro^ dimers bounded to them. The enzyme monomers preferentially bind to the octapeptide, without being released from the it. The octapeptide remains bound to the monomer or dimer (Chou’s “distorted key” theory) [31], [32], [33].

Before proteolytic processing of the viral pp1a and pp1ab into a total of 15 or 16 non-structural proteins (Nsp) occurs, M^pro^ itself is embedded in these polyproteins as the Nsp5 domain. Therefore, the M^pro^ has to first liberate itself from the polyproteins through autocleavage, and then the self-released mature M^pro^ would form a dimer and *trans*-cleave pp1a and pp1ab at other sites. In fact, according to this model,

[6] indicate that N-terminal autocleavage of SARS-CoV Mpro from the polyproteins only requires two “immature” proteases approaching one another to form an “intermediate” dimer structure and does not depend on the active dimer conformation existing in the mature protease that is vital to *trans*-cleavage activity of the protease. In fact, in order to auto-cleave from the replicase polyproteins, the “immature” M^pro^ not yet auto-cleaved can release itself from the polyproteins by inter-molecular cleavage, and then the self-released mature M^pro^ triggers the *trans*-cleavage processing of the other polyproteins. The formation of the “intermediate” dimer could trigger the rotation of their domains I/II relative to domains III and thereby insert their “uncleaved” N-termini into the substrate-binding pockets of the opposite monomer, which might induce the active conformation of the S1 subsites through an induced-fit catalytic mechanism. Once the auto-cleavage of the N-terminus is finished, the “cleaved” N-terminal fingers should slip away from the active sites and switch to their final spatial positions, thereby locking the dimer in a catalytic competitive state. The “uncleaved” C-terminus of one mature dimer can then insert into an active site of another mature dimer and be cleaved. The same occur then to the other mature dimer (this paragraph is an exerpt from [6]).

If almost all the active sites of the “intermediate” dimers are occupied by the synthetic octapeptide substrate inhibitor AVLQSGFR, the “intermediate” monomers cannot cleave each other. As a consequence, the formation of mature M^pro^ will be reduced and the total concentration of the enzyme [*E*]_*T*_ will go down: the higher the concentration of this synthetic peptide in the cell, the lower the total concentration of enzyme in the host’s cells. In this thought experiment, we have indeed reached an interesting theoretical dynamic equilibrium where the catalytic rate of the virus decreases to near zero as we decrease [*E*]_*T*_ by preventing the auto-release of M^pro^. The peak of the curves in Fig. 4 and 5 is then reduced to almost zero and shifted slightly to the left of the graph. Furthermore, in this thought experiment, we dilute the enzyme with the octapeptide substrate, *i*.*e*. we “bathe” the enzyme in a high concentration of (synthetic) substrate. The equilibrium is then further shifted to the right of the curve ([*E*]_*T*_ *≪* [*S*]) (Fig. 4 and 5), with an even lower *V*_*cat*_. Thus, in theory, we are able to control the virulence of the virus by only changing the concentration of the AVLQSGFR octapeptide in the host cell. This mechanism still needs to be verified by experiment. I encourage the scientific community to do so.

## 5 Conclusion

In this study, I developed a model of the reaction mechanism called substrate-induced dimerization. I used the SARS-CoV-2 M^pro^ system as a framework to apply our model *in vitro*. The mathematical development predicts a non-monotonic response of the catalytic rate to the substrate concentration. I have established an equation that describes the behavior of such a system. What seems at first glance to be counterintuitive is the fact that the behavior of the catalytic velocity does not behave like the one of the well-known Michaelis-Menten kinetic model. On the contrary, *V*_*cat*_ shows no maximum asymptote after substrate saturation in our model (*V*_*max*_). Rather, our model behaves similarly to the one describing the substrate inhibition mechanism. In fact, this latter model is quite common in biology [7], [35], [36]. Substrate inhibition is the most common deviation from Michaelis-Menten kinetics, occurring in about 25% of known enzymes. It is generally attributed to the formation of an unproductive enzyme-substrate complex after the simultaneous binding of two or more substrate molecules to the active site [37]. Thus, our model behaves much as in the case of a substrate inhibition mechanism, with one important exception: in our case, we have two enzyme monomers bound to one substrate molecule, whereas the substrate inhibition model considers the *ES*_2_ system, *i*.*e*. two substrate molecules bound to one enzyme molecule, which is actually a radically different type of model from ours. The only element that brings the two models together is the non-monotonic curve, which is present in both models.

Thus, I propose a new mathematical description of substrate-induced dimerization, which we encounter in the SARS-CoV-2 M^pro^ system. I show that the behavior of this enzyme(s)-substrate system implies that at high substrate concentration, we have only weak enzymatic kinetics due to the dilution of the enzyme monomers in the solution saturated with substrate molecules (see Fig. 2). In other words, this low enzyme activity means that the amount of substrate in solution determines the activity of the key enzyme M^pro^, and thus probably the virulence of the coronavirus. This means that we could theoretically take advantage of this phenomenon by inducing a high concentration of substrate, which thus leads to a decrease in the activity of M^pro^, or, as it were, of the pathogen. Moreover, I found *V*_*max*_ and 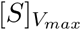 analytically from the mathematical model. I compared these expressions with the different fitted curves, and found that the numerical values correspond very well to the practical values of the fitted curves. Finally, I may be able to propose a type of substrate that would work as M^pro^-diluting factor. The synthetic octapeptide substrate AVLQSGFR functions as a specific inhibitor of M^pro^ (either in the SARS-CoV-2 or SARS-CoV system, which are biologically very similar). I have deduced that this octapeptide can potentially and theoretically shift the peak of the curves of Fig. 4 and 5 by reducing the concentration of the total enzyme in the host’s cell. This can be done by docking the octapeptide directly to the “intermediate” dimers at their active sites.

## 6 Outlook

In this outlook, I want to try to generalize the model I presented above and apply it to animal ecology. This is only an attempt without “proof of principle”. The discussion below serves only to broaden the spectrum of knowledge discussed in this study. There is no proof or claim that this new concept described below also works in reality. Nevertheless, I wanted to present it here from this perspective.

An attempt has been made to synthesize the generalized model of plant nutrient harvest rate with predator foraging theory by considering prey harvest rate. In this regard, one approach proposed that the single-species Holling’s disc equation used by animal ecologists to model resource uptake ([40], [41], [42]) and the Michaelis-Menten equation used by plant physiologists to model nutrient uptake ([1], [43]) are actually rearranged forms of the same functional response [44] (this latter is defined as the number of preys consumed per predator as a function of prey density [48]). Indeed, a graphical comparison of the Michaelis-Menten equation and the Holling’s disc equation shows two very similar curves: these two descriptions are mathematically equivalent [50] [44]. Bearing this in mind, we can deduce that this type of generalization could also be applied to our model described above. In order to extrapolate equations (19) to (24) and the graph modeling the SARS-CoV-2 M^pro^ system to a completely different system (a predator-prey model in animal ecology for example), we need to find another animal ecological model that is broadly compatible with our theory of enzymatic kinetics (*i*.*e*. substrate-induced dimerization of SARS-CoV-2 M^pro^). But, in this second system, which we will now call the “predator-prey model”, what is the monomeric and/or dimeric enzyme counterpart, and what is the substrate counterpart? I will try to explain my reasoning and put forward my arguments.

After a thorough search in literature, I found the following statement: cooperative hunting is probably the most widely spread form of cooperative behavior in animals. The Aplomado (*Falco femoralis*) [52], Lanner (*Falco biarmicus*) [53], and Saker (*Falco cherrug*) falcons [55], for example, or the Bonelli’s eagle (*Hieraaetus fasciatus*) and sometimes the Golden eagle (*Aquila chrysaetos*), hunt in pairs, assuming that the incubation period has occurred, the chicks have fledged, and the female has joined the male in hunting. In the latter case, when hunting in pairs, the male Golden eagle, for example, flies in front of the female at a higher altitude and usually takes the initiative to attack. The first pursuer diverts the attention of the prey by ducking while the second flies away unseen to shoot it down (see [57] and [58]). This means that, in this case for example, the male and female are hunting the same prey in tandem. Hector (1986) [52] has shown that Aplomado falcons regularly hunt in pairs with successful tandem hunts of avian prey 45% of the time versus 21% for solo hunts. Thiollay (1988) [45] reported that 37.8% of peregrine falcon (*Falco peregrinus*) pair incursions were successful, compared to 16.5% for males and 23.3% for females hunting solo. Further on, cooperative hunting may provide greater protection to each individual against larger kleptoparasitic raptors. In addition, the higher success rates of cooperative hunts may reduce the time required to provision youngs and leave more time for nest defense [52].

Predator-prey relationships are the result of an evolutionary arms race in which the prey adopts strategies to reduce the risk of being captured [46]. I present here an excerpt from [52], p. 252: “*During spring migration, falcons attacked migrating flocks of white-winged doves, and mourning doves. The falcons would ascend rapidly from lookout posts then fly directly at flocks still 3-4 km away and 50-200 m above the ground. In most cases, dove flocks did not alter course and scatter until falcons closed to within 50 m. Both falcons then pursued the same individual on a slanting or vertically descending path. Often these chases would level off near the ground with the female following immediately behind the dove, and the male flying overhead to begin a series of dives and ascents*.”

It has been shown that in some cases the functional response to a single resource exhibits a Type IV dome-like appearance due to prey overabundance, *i*.*e*. a non-monotonic curve with only one peak (*i*.*e*. unimodal). The Type IV response is the only functional response that does not increase monotonically with increasing resource density. Until now, the Type IV response has been described by an equation similar to the Type II Michaelis–Menten equation, but with additional term in the denominator (*βN* ^2^, with “N” corresponding to the prey density and *β* a constant) [59]. In ecology, the dome-shaped functional response can be explained, for example, by the phenomenon of prey toxicity [59], or predator confusion [60]. This latter case is an explanation of the “dilution” effect: for any attack by a predator, the larger the group of prey, the lower the probability that a particular individual will be the victim (swarm or flock effect). Prey-grouping may make it difficult for a predator to single out prey [61], [62].

Let me now argue as follows. Assuming that we are able to draw parallels between the monomeric SARS-CoV-2 M^pro^ enzyme and a predatory individual on the one hand, and between the TQ6-pNA substrate and a prey individual on the other, we could imagine that the two kinds of model have some commonalities: in this regard, the substrate-induced dimerization process could be extrapolated to an analogous process, but adapted to the raptor-prey model. In our case, eqs. (19) to (24) are a putative alternative description of the dome-shaped Type IV functional response, which could be suitable to describe certain types of predator-prey interactions. In this regard, the sequential binding of two monomeric M^pro^ enzymes to the TQ6-pNA substrate is actually comparable to the hunting scenario of two Golden eagles focusing on a single prey. In this theoretical and simplified model, the first of the two eagles needs the help and assistance of its partner to kill the prey. This behavior has the same mechanistic basis as the one of two enzymes that must be bound together to the same substrate to be active and degrade it. In both models, we need two identical components (*i*.*e*., the two M^pro^ monomers, respectively the two raptor individuals), and one substrate molecule (TQ6-pNA), resp. one prey individual (e.g. a dove or starling), where the two identical components need to be docked (to the substrate resp. to the prey) in order to degrade enzymatically (the substrate) resp. kill (the prey).

Also, we assume that our new raptor-prey model has to match the non-monotonic curve shape of our original SARS-CoV-2 M^pro^ enzymatic model. As described in the second paragraph of this section, we suppose that the functional response curve of the raptor-prey model has the same appearance as the curve derived from our calculations above. This latter curve has also a dome-shaped appearance and is also unimodal. We see in nature that raptors hunt preys that are sometimes “diluted” in a flock. Predators may have difficulty catching and killing their prey because of the “dilution” of the preys in the swarm and the resulting confusion of the predator confronted with the swarm of its prey. The predator cannot focus as well on a single prey, and if it does, the second predator is not very able to dock on the resulting first predator-prey pair because of this confusion state. In fact, this phenomenon corresponds to the same pattern detailed in our study: in part C of Fig. 2, the monomers of the enzyme are “diluted” in the solution, the probability of having two monomers of the enzyme docked on the same substrate molecule is low. Indeed, the more substrate or prey entities are present, the more diluted are the enzyme monomers in the solution, resp. the raptor individuals in the swarm. Ideally, for [*S*] *≫* [*E*]_*T*_ or [prey] *≫* [predator], we have *V*_*cat*_ or *I* (“*I*” being the prey intake) tending to 0 when [*S*] or [prey] tend to *∞*. On the contrary, in part B of Fig. 2 above, at a defined [*S*]_*max*_ resp. [prey]_*max*_ (*i*.*e*. the density of substrate or prey for which the catalytical rate or the number of prey consumed are maximum), there is a maximum of interactions between the enzyme monomers / raptor individuals and the substrate molecules / prey individuals for which the number of prey consumed is maximal (top of the dome-shaped curve). The efficacy of the substrate-induced dimerization/tandem-raptor attacks is thus at its maximum here. In this case, we have [*E*]_*T*_ ≈ 2[*S*]_*max*_ or [predator] ≈ 2[prey], where the symbol [ ] represents the density in this latter case. Finally, in part A of Fig. 2, we have the case where [predator] *≫* [prey], or equivalently, [*E*]_*T*_ *≫* [*S*]. There is here an excess of predator entities, so it is not very difficult for a predator pair to focus on a prey and, after attacking it, kill it. Conversely, the enzyme monomers are in excess in solution in this case, and two monomers can successfully sequentially dock to a substrate molecule by the mechanism of substrate-induced dimerization. In this case, we consider the part to the left of the maximum of the dome-shaped functional response, which is hyperbolic and similar to the Michaelis-Menten curve. The parallelism is striking between the substrate-induced dimerization model and the predator-prey model explained here (with the above given assumptions).

I want to make it clear that these are only theoretical considerations, and that this raptor-prey model I present here still needs to be validated by experiment. One of the main pitfalls of this model compared to the substrate-induced dimerization model explained above is that in this former model we are dealing with living entities, e.g., with doves forming a flock that evolves strongly and in a coherent way in space over time, certainly more than substrate molecules in solution which are more static and disorganized in comparison. When starling (*Sturnus vulgaris*) flocks are under attack by a raptor, such as a Peregrine falcon, they show a great diversity of patterns of collective escape [64]. The corresponding structural complexity concerns rapid variation in density and shape of the flock over time. I acknowledge that my reasoning corresponds to a strongly idealized model, which is perhaps only valid in a very limited way. I would like to mention this raptor-prey model in order to show that this new enzymatic M^pro^ model could possibly be at the source of an extrapolation in the field of animal ecology.

To conclude, I would like to take as an example an article describing the use of a pair of falcons against doves in the city of Tournus in France. This article [65] mentions that the doves are not killed, but only frightened. This is consistent with our prediction that, when the prey behaves in a swarm, the raptor has difficulty to kill it, in solo or especially in tandem.

## 7 Acknowledgements

I thank Felix Kessler for inviting me in his lab as a visiting scientist. I am grateful to Marcelle Kaufman and Didier Gonze for their fruitful commentaries on section 3 of the manuscript. I thank my family for their support.

## 9 Additional information

Correspondence and requests for materials should be addressed to Thierry Rebetez.

### Open Access

This article is licensed under a Creative Commons Attribution 4.0 International License, which permits use, sharing, adaptation, distribution and reproduction in any medium or format, as long as you give appropriate credit to the original author and the source, provide a link to the Creative Commons license, and indicate if changes were made. The images or other third party material in this article are included in the article’s Creative Commons license, unless indicated otherwise in a credit line to the material. If material is not included in the article’s Creative Commons license and your intended use is not permitted by statutory regulation or exceeds the permitted use, you will need to obtain permission directly from the copyright holder. To view a copy of this license, visit http://creativecommons.org/licenses/by/4.0/.

